# Islet transplantation tolerance in animals with defined histocompatibility and diabetes

**DOI:** 10.1101/2021.10.08.463702

**Authors:** Preksha Bhagchandani, Charles A. Chang, Weichen Zhao, Luiza Ghila, Pedro L. Herrera, Simona Chera, Seung K. Kim

**Affiliations:** Department of Developmental Biology, Stanford University School of Medicine, Stanford, CA, 94305, USA; Department of Clinical Science, University of Bergen, Bergen, Norway; Department of Genetic Medicine and Development, University of Geneva, Geneva, Switzerland; Department of Medicine (Endocrinology Division), Stanford University School of Medicine, Stanford, CA 94305, USA; Department of Pediatrics (Endocrinology Division), Stanford University School of Medicine, Stanford, CA 94305, USA; Stanford Diabetes Research Center, Stanford University School of Medicine, Stanford, CA, 94305, USA; JDRF Center of Excellence, Stanford University School of Medicine, Stanford, CA, 94305, USA

## Abstract

Advances in organ transplantation benefit from development of genetically inbred animal strains with defined histocompatibility and cell-specific markers to distinguish donor and host cell subsets. For studies of pancreatic islet transplantation tolerance in diabetes, an invariant method to ablate host β cells and induce diabetes would provide an immense additional advantage. Here we detail development and use of *B6 RIP-DTR* mice, an immunocompetent line permitting diabetes induction with 100% penetrance. This inbred line is homozygous for the C57BL/6J major histocompatibility complex (MHC) haplotype and expresses the mutant *CD45.1* allele in the hematopoietic lineage. β cell-specific expression of a high-affinity receptor for diphtheria toxin (DT) permits experimental β cell ablation and diabetes induction after DT administration. Diabetes reversal for over one year was achieved after transplantation with congenic C57BL/6J islets, but not with MHC-mismatched BALB/c islets, which were rapidly rejected. In summary, the generation of a C57BL/6J congenic line harboring the *CD45.1* allele and *Ins2-HBEGF* transgene should advance studies of islet transplantation tolerance and mechanisms to improve islet engraftment and function, thereby optimizing development of cell replacement strategies for diabetes mellitus.

## Introduction

Studies of transplantation tolerance in mice have relied on generation of inbred strains with defined histocompatibility.^1^ For example, inbred mouse lines homozygous for defined major histocompatibility complex (MHC) loci have been used to improve immunomodulation, develop targeted anti-inflammatory strategies, and identify non-myeloablative bone marrow conditioning regimens to achieve mixed hematopoietic chimerism.^2^ For islet transplantation studies, robust genetic methods for reliable diabetes induction in immunocompetent transplant recipients could accelerate advances in islet allotolerance.

In prior studies^3^, we generated mice permitting genetically-encoded β cell ablation and robust diabetes induction. Pancreatic islet β cells in these mice express the membrane-anchored form of human heparin binding EGF-like growth factor (HB-EGF), which binds diphtheria toxin (and is hereafter referred to as ‘diphtheria toxin receptor’; DTR). Upon diphtheria toxin (DT) administration, β cells expressing DTR are unable to synthesize protein due to inhibition of elongation factor 2 (EF2), an essential protein for ribosomal translocation, leading to cell death.^4^ Targeted HB-EGF expression in β cells is directed from a rat insulin promoter (RIP) element, *Ins2*. Since mouse and rat cells do not bind DT, DTR-mediated cell ablation is a highly sensitive and specific method that allows targeting of β cell in RIP-DTR mice without off-target host cell damage or toxicity.^5^ In RIP-DTR adult mice, DT injection results in >99% β cell ablation, with low overall β cell regeneration rate thereafter.^3^ Work in islet transplantation biology and immunology would benefit from development of inbred RIP-DTR mice with defined histocompatibility.

Here we detail development and use of an inbred, immunocompetent strain harboring the RIP-DTR transgene, which permits diabetes induction via DTR-mediated β cell ablation with 100% penetrance. This strain is hereafter called “B6 RIP-DTR.”

## Results

### Generation, genetics, and immunological phenotypes of B6 RIP-DTR mice

The original RIP-DTR strain was maintained on a genetically ‘mixed’ B6/CBA background.^3,6,7^ To make RIP-DTR mice suitable for organ transplantation studies, we intercrossed these repeatedly (‘backcrossed’) to B6.SJL-*Ptprc*^*a*^ *Pepc*^*b*^/BoyJ mice, commonly referred to as Pep Boy or B6 CD45.1 (hereafter, B6 CD45.1), to generate an inbred line with a defined histocompatibility genotype. After backcrossing, we analyzed B6 CD45.1 RIP-DTR mice (abbreviated B6 RIP-DTR) using single nucleotide polymorphism (SNP) genome scanning analysis (**Table 1**) to measure the % C57BL/6J identity. After multiple backcrosses to B6 CD45.1 (**Methods**), we achieved an average 96% C57BL/6J identity at 120 single nucleotide polymorphisms (SNPs) analyzed across 20 chromosomes from 12 mice. Most notably, we achieved 100% C57BL/6J identity on chromosome 17, which contains all class I and class II major histocompatibility complex (MHC) genes responsible for antigen presentation and distinguishing self from non-self.^8^ Thus, our genotyping indicated successful generation of inbred B6 RIP-DTR mice.

**Table 1:** Analysis of percentage CBA/J and C57BL/6J background in RIP-DTR mice (n = 12, 6F, 6M) following final backcross to B6 CD45.1 background. Chromosome 17 contains Major Histocompatibility Complex (MHC) genes. SNP = single nucleotide polymorphism

SNP Genome Scanning Analysis of CBA and B6 Backgrounds
| *Chromosome* | *Number of*<br>*SNPs*<br>*assessed* | *Average*<br>*Percentage*<br>*C57BL/6J* | *Average*<br>*Percentage*<br>*CBA/J* |
| --- | --- | --- | --- |
| 1 | 9 | 70% | 30% |
| 2 | 9 | 100% | 0% |
| 3 | 5 | 92% | 8% |
| 4 | 8 | 100% | 0% |
| 5 | 6 | 96% | 4% |
| 6 | 7 | 100% | 0% |
| 7 | 7 | 94% | 6% |
| 8 | 6 | 97% | 3% |
| 9 | 5 | 100% | 0% |
| 10 | 6 | 100% | 0% |
| 11 | 6 | 98% | 2% |
| 12 | 6 | 93% | 7% |
| 13 | 5 | 100% | 0% |
| 14 | 5 | 100% | 0% |
| 15 | 5 | 100% | 0% |
| 16 | 5 | 100% | 0% |
| 17 | 5 | 100% | 0% |
| 18 | 5 | 100% | 0% |
| 19 | 4 | 100% | 0% |
| X | 6 | 84% | 16% |
| **Overall** | **120** | **96%** | **4%** |

MHC genes are inherited together as they are clustered on mouse chromosome 17 and closely linked.^9^ To phenotype the MHC haplotypes of B6 RIP-DTR mice, we used flow cytometry to analyze peripheral blood lymphocyte marker production (**Figure 1, Table 2**). The *MHC Class II* gene I-A is expressed on antigen-presenting cells, and the I-A^b^ allele is expressed by C57BL/6J mice, while the I-A^k^ allele is expressed by CBA/J mice. I-A^b^ was expressed by blood cells in both C57BL/6J and B6 RIP-DTR mice, while I-A^k^ was expressed in CBA/J mice (**Figure 1A**). I-E^k^, another *MHC Class II* gene, was expressed by CBA/J blood but was not detectable in C57BL/6J nor B6 RIP-DTR mice (**Figure 1B**). The *MHC Class I* gene H-2K is expressed ubiquitously, and the H-2K^b^ allele is characteristic of C57BL/6J mice while H-2K^k^ is characteristic of CBA/J mice. Flow cytometry revealed that H-2K^b^, not H-2K^k^, was produced by blood cells in both C57BL/6J and B6 RIP-DTR mice (**Figure 1C**). Thus, molecular phenotyping confirmed that B6 RIP-DTR mice express *MHC Class I* and *Class II* gene products characteristic of C57BL/6J mice.

**Table 2:** MHC haplotypes for C57BL/6J, CBA/J, and BALB/cJ mice

MHC haplotypes for C57BL/6J, CBA/J, and BALB/cJ mice
| Mouse Strain | MHC Haplotype | *MHC Class I* |  |  | *MHC Class II* |  |
| --- | --- | --- | --- | --- | --- | --- |
|  |  | H-2K | H-2D | H-2L | I-A | I-E |
| *C57BL/6J* | b | b | b | null | b | null |
| *CBA/J* | k | k | k | null | k | k |
| *BALB/cJ* | d | d | d | d | d | d |

**Figure 1:**
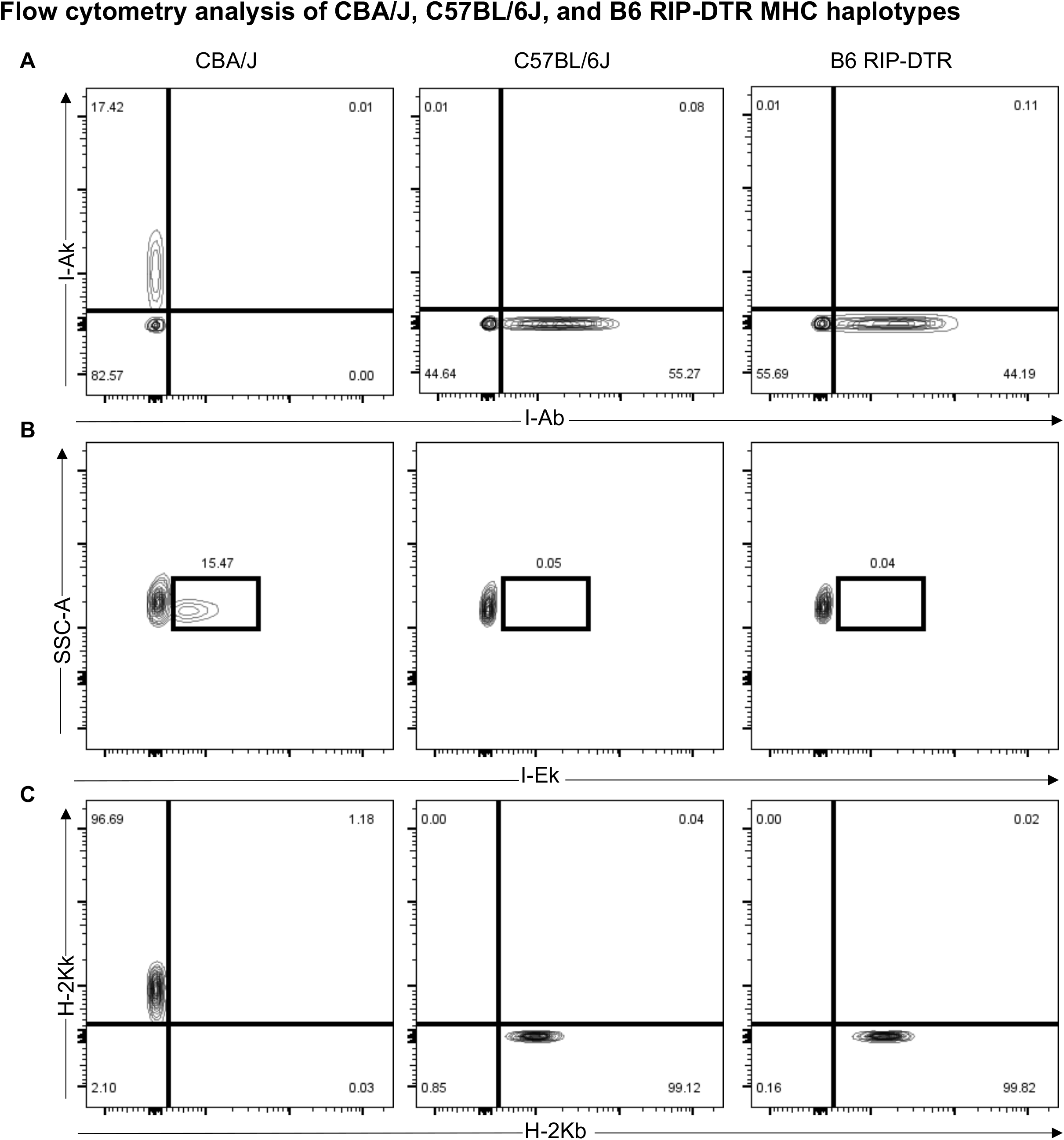
**A)** MHC Class II alleles I-A^b^ (C57BL/6J haplotype), I-A^k^ (CBA/J haplotype) and **B)** I-E^k^ (CBA/J haplotype) analyzed on C57BL/6J and CBA/J controls, as well as B6 RIP-DTR mice. **C)** MHC Class I alleles H-2K^b^ (C57BL/6J haplotype), H-2K^k^ (CBA/J haplotype) analyzed on C57BL/6J and CBA/J controls, as well as B6 RIP-DTR mice.

### RIP-DTR and CD45.1 alleles in B6 RIP-DTR mice

To empower future studies in B6 RIP-DTR mice receiving allogeneic hematopoietic cell transplantation, we sought to make immune cells in B6 RIP-DTR mice readily distinguishable from allogeneic donors. Most available donor strains express the CD45.2 epitope on immune cells. Thus, our breeding strategy aimed to produce B6 RIP-DTR homozygous for the mutant CD45.1 allele, which encodes a cell surface protein readily distinguishable from CD45.2 using flow cytometry.^10^ To achieve this goal, we backcrossed RIP-DTR mice with B6 CD45.1 mice (**Figure 2**). We verified the presence of the RIP-DTR transgene using PCR (**Figure 2A; Methods**). We also generated genotyping tools (**Methods**) to track and confirm homozygosity of the CD45.1 allele. Specifically, dual endpoint qPCR was used to distinguish the CD45.1 allele from CD45.2, in the initial backcrosses and all subsequent generations of B6 RIP-DTR CD45.1 mice (**Figure 2B**). These approaches generated inbred B6 RIP-DTR mice homozygous for the CD45.1 allele.

**Figure 2:**
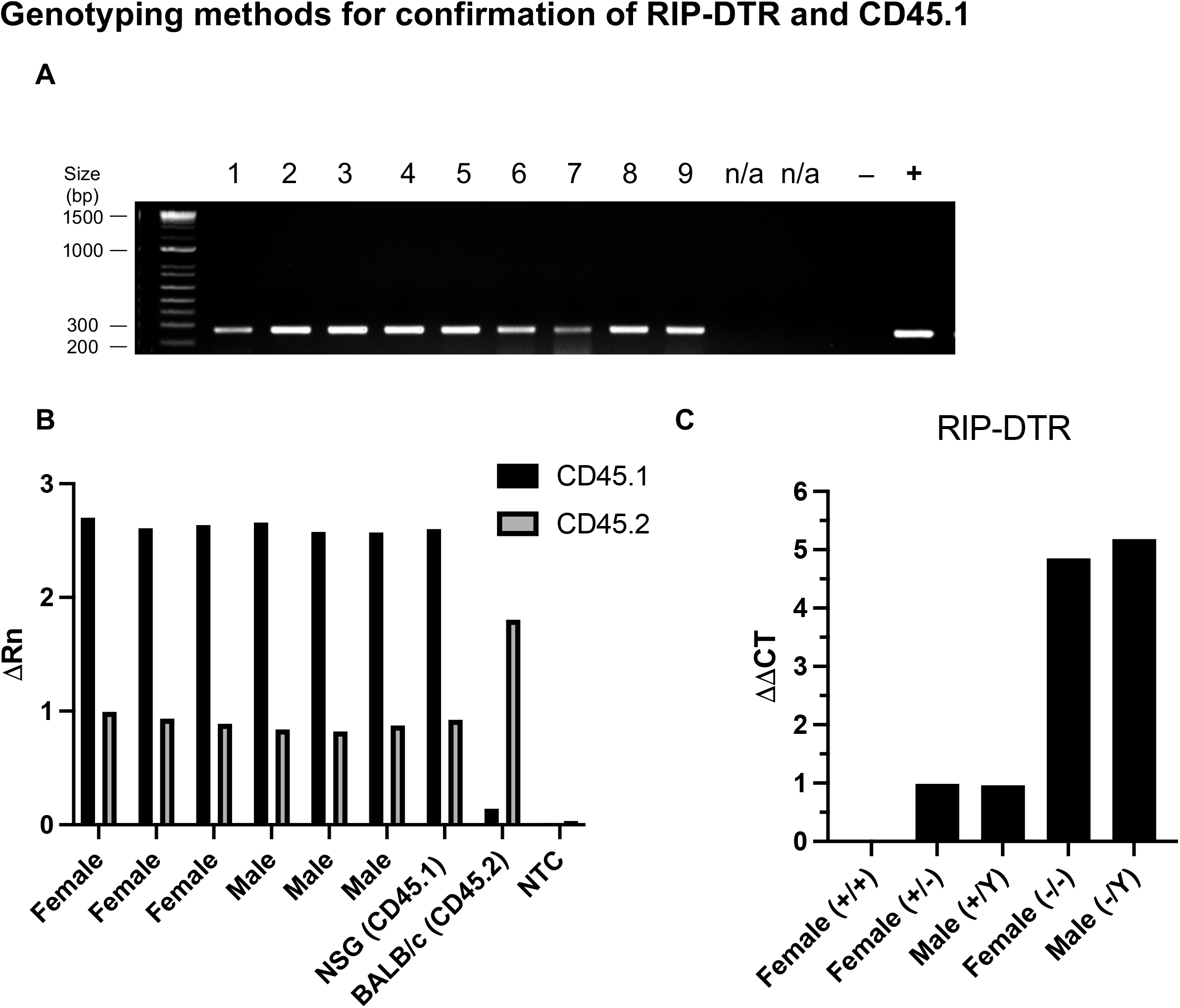
**A)** Agarose gel electrophoresis (2% agarose) of PCR amplified products using RIP-DTR specific PCR primers. Lanes 1-9 are RIP-DTR males and females. Negative control is a B6 mouse. Positive control is a mouse that was confirmed to have RIP-DTR by DT injection. Bp = base pairs. **B)** Dual endpoint qPCR analysis for CD45.2 or CD45.1 allele of CD45 in female and male RIP-DTR mice (n = 6). NSG and BALB/c mice were used as controls for CD45.1 and CD45.2, respectively. NTC = No Template Control. Rn = normalized reporter value **C)** Comparative qPCR of RIP-DTR compared to CD45 to determine zygosity of RIP-DTR females. (+/+) denotes homozygous, (+/-) denotes heterozygous, and (+/Y) denotes hemizygous for RIP-DTR, (-/-) and (-/Y) denote absence of RIP-DTR. CT = cycle threshold

Since the RIP-DTR transgene is X-linked, female RIP-DTR heterozygotes remain normoglycemic after diphtheria toxin administration due to random X chromosome inactivation in β cells.^3^ Thus, we sought to produce female B6 RIP-DTR homozygous for RIP-DTR, and males hemizygous for RIP-DTR. To achieve this, we developed a comparative qPCR method to distinguish female heterozygotes from homozygotes (**Figure 2C**). Homozygous females (2 copies of RIP-DTR) result in a one cycle threshold (CT) difference from heterozygous females with one copy of RIP-DTR. Thus, our methods reliably measured RIP-DTR copy number, and generated breeder pairs that ensure production of B6 RIP-DTR female mice homozygous for RIP-DTR and males hemizygous for RIP-DTR. All experiments were performed on mice with these genotypes.

### Overt diabetes in RIP-DTR mice after diphtheria toxin injection

To test the efficiency of diabetes induction in our colony of RIP-DTR mice, we administered a single dose of diphtheria toxin (DT) intraperitoneally (i.p.) to males (n = 16) and females (n=13) between 8 and 24 weeks of age (**Figure 3; Methods**). Blood glucose was measured before and after injection until mice were overtly hyperglycemic (>250 mg/dL). 100% of male mice were severely hyperglycemic approximately 3 days after DT injection, regardless of age (**Figure 3A**). Likewise, 100% of female mice were severely hyperglycemic approximately 4 days after DT injection, regardless of age (**Figure 3B**). Within 24-48 hours after the onset of hyperglycemia, the health of diabetic mice deteriorated, reflected by rapid weight loss, unless provided exogenous insulin. The observed sex difference between male and females for time to induction could reflect differences in sex hormones^11,12^. Regardless, we were able to predict the timing of hyperglycemia onset for both sexes for subsequent islet transplantation studies. Additionally, histology of pancreas taken from RIP-DTR mice at 4 months after single DT injection shows effective β cell ablation, with little to no insulin producing cells remaining (**Figure 3C**). By contrast, histology of the pancreas from control B6 CD45.1 mice injected with DT revealed normal-appearing islets that included intact β cells and other islet cells (**Figure 3D**). Thus, our studies revealed that the RIP-DTR transgene was functional in B6 RIP-DTR mice, and results in rapid and reliable diabetes induction.

**Figure 3:**
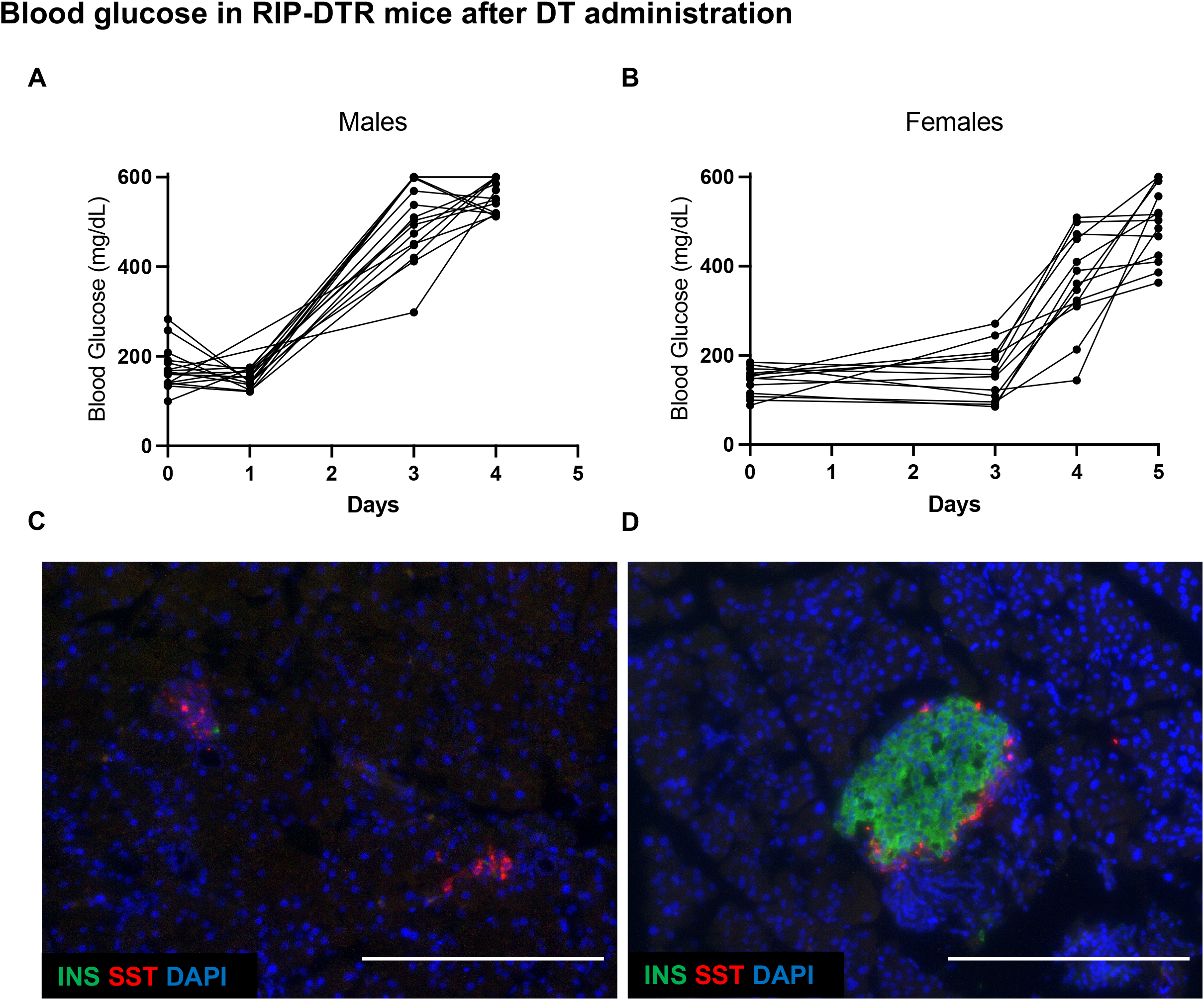
Non-fasting blood glucose in RIP-DTR mice after single dose administration of DT (i.p.) in n = 16 hemizygous males (**A)** and n=13 homozygous females (**B**). DT was injected on day 0 after baseline non-fasting blood glucose was recorded. Mice were between 8 and 24 weeks of age at time of injection. Representative histology of pancreas taken at 4 months from B6 RIP-DTR (**C**) or B6 CD45.1 (**D**) mice given single dose of DT. INS = Insulin. SST = somatostatin. Scale bar = 200 µm.

### Islet transplantation tolerance and allorejection in B6 RIP-DTR mice

Similar to the commonly used C57BL/6J mouse strain, B6 RIP-DTR have multiple phenotypes suitable for islet transplantation studies. This included expected average lifespan, litter sizes, and breeding performance (**Methods**). For example, expected body mass at 10 weeks of age was similar in B6 RIP-DTR and C57BL/6J mice. In sum, the physiology, breeding and growth characteristics of B6 RIP-DTR make them well-suited for islet transplantation studies.

To assess the use of B6 RIP-DTR mice for studies of immune tolerance, we transplanted these mice with congenic B6 islets or BALB/c islets, which are fully MHC-mismatched (**Figure 4A**). Three B6 RIP-DTR mice, at 10 weeks of age, were injected with DT, and after hyperglycemia onset were transplanted in the renal sub-capsular space with islets from C57BL/6J donors. By three days after transplantation, all recipients became euglycemic without supplemental insulin and have remained so for at least 56 weeks following islet transplantation (**Figure 4B**). Reversion to hyperglycemia invariably followed nephrectomy and islet graft removal at 56 weeks (**Figure 4B**); thus, diabetes reversal in these B6 RIP-DTR mice prior to nephrectomy reflected the function of transplanted islets. In summary, congenic C57BL/6J islets are durably tolerated and functional after transplantation in the novel B6 RIP-DTR mouse strain.

**Figure 4:**
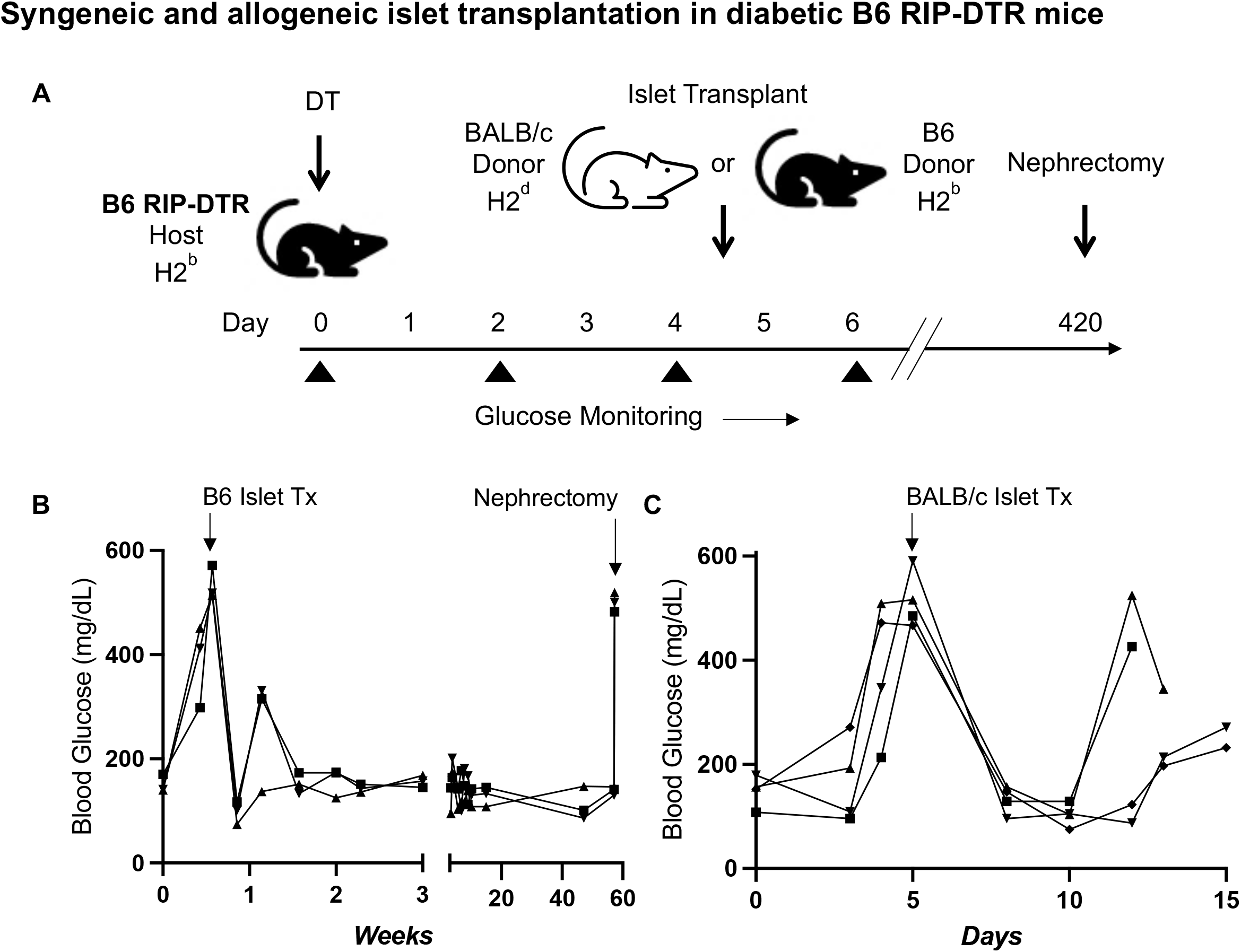
**A)** Schematic of congenic and allogeneic islet transplantation in diabetic B6 RIP-DTR mice. **B)** Male RIP-DTR mice (n = 3; age 10 weeks) were injected with DT on day 0 and transplanted with congenic C57B6/J islets on day 4 after confirmation of hyperglycemia. Mice were monitored for approximately 1 year, until nephrectomy was performed at 56 weeks post-transplant to remove the islet graft. **C)** Female RIP-DTR mice (n = 4; age 10 weeks) were injected with DT on day 0 and transplanted with BALB/c (allogeneic) islets on day 5 after confirmation of hyperglycemia. Mice were monitored for rejection until confirmation of repeat hyperglycemia.

To assess the immunocompetence of B6 RIP-DTR mice for rejecting islets from mismatched BALB/c donors (**Table 2**), we transplanted BALB/c islets into four B6 RIP-DTR mice after administration of DT and confirmation of hyperglycemia (**Figure 4C**). After transplantation, all mice became euglycemic, demonstrating effective engraftment and functional status of the transplanted islets. However, within two weeks of transplantation, all four mice had reverted to overt hyperglycemia, consistent with the timeline of adaptive immunological rejection. These results demonstrate that B6 RIP-DTR mice remain immunocompetent for rejecting allogeneic donor islets.

## Discussion

The RIP-DTR mouse model of inducible diabetes has proven useful for studies of conditional beta cell loss and regeneration.^3,7^ β cell ablation in mice harboring the RIP-DTR transgene is reproducible with 100% penetrance, and more than 99% complete.^3^ However, the mixed B6/CBA genetic strain background of the original RIP-DTR mice precluded studies of allotolerance, specifically in islet transplantation. Here we detail use of intercrosses, genotyping and molecular phenotyping to produce a genetically inbred strain of B6 RIP-DTR mice, and use of this new strain for studies of conditional β cell ablation and diabetes, islet allotolerance and rejection. Moreover, homozygous incorporation of the CD45.1 allele should permit powerful flow cytometry-based studies of transplanted hematopoietic cell lineages^13^, including immune cell subsets, in B6 RIP-DTR mice.

The RIP-DTR transgene has been used to model β cell death, without immune infiltrates.^3^ Diphtheria toxin administration initiates cell lysis and intranucleosomal DNA fragmentation (apoptosis); since mouse cells lack the DT receptor, cells in transgenic mouse strains expressing a human transgene encoding this receptor can be selectively destroyed.^14^ In contrast to this targeted effect, chemotoxic methods to destroy β cells, like streptozotocin (STZ) challenge^10,15,16^, often produce unwanted damage and off-target effects in liver, nervous system, cardiovascular system, respiratory system, kidneys, and reproductive system.^10^ Other limitations of STZ challenge include variability of responses in different strains and sexes.^10,15,16^ Development of tools here to ensure the RIP-DTR genotype and highly penetrant DT-dependent β cell ablation in all B6 RIP-DTR progeny should promote broader use of this strain.

Diabetes reversal after transplantation with congenic B6 islets but not with MHC-mismatched BALB/c islets provides an index demonstration of the uses of in B6 RIP-DTR mice for islet allotolerance and rejection studies. In summary, the unique combination of C57BL/6J congenic MHC haplotype, *CD45.1* mutant allele, and the *Ins2-HBEGF* allele in the B6 RIP-DTR strain should advance studies of islet transplantation tolerance, including investigation of mechanisms to improve islet engraftment and function, like hematopoietic chimerism to promote islet allotolerance in the setting of diabetes.

## Material and Methods

### Animals

Female and male B6 CD45.1 (Stock #: 002014), BALB/c (Stock #: 000651), CBA (Stock #: 000656), and NSG (Stock #: 005557) mice were purchased from The Jackson Laboratory (Bar Harbor, ME). RIP-DTR mice were originally on a mixed B6/CBA genetic background and maintained by Dr. Simona Chera at UIB before transferring to Dr. Seung Kim at Stanford. Female mice were backcrossed with male B6 CD45.1 for multiple generations until >95% B6 background and homozygosity for CD45.1 were achieved. SNP genome scanning analysis was performed by sending samples to The Jackson Laboratory (Bar Harbor, ME) after each backcross. Sibling matings with verified heterozygous offspring were used to further breeding to RIP-DTR homozygosity after each backcross. B6 RIP-DTR mice have lifespan of at least one year, and average litter size of 7 +/- 2 (n=9). At 10 weeks of age, average male weight is 27.3 g +/- 2.3 (n = 13), and average female weight is 19.7 g +/- 1.5 (n=13). All animals were housed in non-barrier conditions at the Stanford School of Medicine. Animal experiments were approved by the Stanford Administrative Panel on Laboratory Animal Care.

### Mouse genotyping

Genotyping for the RIP-DTR transgene was performed using PCR amplification with forward primer (GGT GCT GAA GCT CTT TCT GG) and reverse primer (CTC CTC CTT GTT TGG TGT GG), which produce a 250 bp product. PCR products were analyzed by agarose gel electrophoresis. Dual endpoint qPCR was used for CD45.1/2 genotyping. DNA was extracted from tail snips and qPCR was performed with TaqMan Gene Expression Assay kit (Thermo Fisher Scientific, Waltham, MA) according to manufacturer instructions, CD45 forward primer (CGC CTA AGC CTA GTT GTG G) and reverse primer (ATT CTT GAT TTT GTT TCC CTA GTG G), as well as CD45.1 (MUT) probe (CCT GAG CCT GCA TCT AAA CCT G) and CD45.2 (WT) probe (CCT GAG CCT GTA TCT AAA CCT GA). To distinguish homozygous RIP-DTR females from heterozygous RIP-DTR females, comparative qPCR was used. DNA was extracted from tail snips above was cleaned using Genomic DNA Clean and Concentrator kit (Zymo Research, Irvine, CA). qPCR was performed with CD45 and RIP-DTR forward and reverse primers used above and SYBR green master mix (Bimake, Houston, TX). Number of copies of RIP-DTR present was calculated by normalizing RIP-DTR cycle threshold (CT) to CD45 cycle threshold to obtain ΔCT, and again normalizing to a homozygous RIP-DTR female confirmed by DT injection to obtain ΔΔCT. Female homozygotes and heterozygotes exhibited 1 CT difference.

### Flow cytometry analysis of peripheral blood

100 µl of whole blood was collected via the tail vain into EDTA coated tubes. Whole blood was lysed in 1x RBC Lysis Buffer (Biolegend, San Diego, CA) for 10 min on ice before downstream staining. For analysis of MHC alleles, cells were first stained in PBS to distinguish between live and dead cells using LIVE/DEAD Fixable Near-IR Dead Cell Stain Kit (ThermoFisher Scientific). Cells were then incubated with TruStain FcX anti-mouse CD16/32 Fc block (Biolegend) for 10 min on ice in Cell Stain Buffer (Biolegend). Antibodies used for staining from Biolegend were as follows: I-Ak PE (10-3.6), I-Ab AF647 (KH74), H-2Kk FITC (36-7-5), H-2Kb PerCP/Cy5.5 B (AF6-88.5). I-Ek VioBlue (REA510) was purchased from Miltenyi Biotec (Bergisch Gladbach, Germany). Cells were analyzed with a BD FACSAria II. Data were analyzed using FlowJo (10.7).

### Induction of Diabetes, Islet Isolation, and Islet Transplantation

A one-time 500 ng injection of diphtheria toxin (DT) (Cayman Chemical, Ann Arbor, MI) in Hank’s Buffered Salt Solution (HBS; Caisson Labs, Smithfield, UT) was administered intraperitoneally (i.p.) in adult mice to induce diabetes from β cell ablation. Islet isolation and transplantation were performed as previously described with minor modifications.^17,18^ Briefly, pancreases are perfused with 100-125μg/mL Liberase TL (Roche Diagnostics, Indianapolis, IN) and digested in a 37°C water bath for 18-22 minutes. After washing with HBS the crude digest is purified over a discontinuous density gradient, washed once more with HBS, and cultured overnight in RPMI 1640 (Corning; Corning NY) supplemented with 10% FBS, 10mM HEPES, and 1% penicillin-streptomycin solution. Recipient mice are anesthetized with ketamine and xylazine and given subcutaneous analgesics. After overnight culture, 100-400 islets are loaded into polyethylene (PE)-50 tubing (BD, Franklin Lakes, NJ) and injected under the kidney capsule of recipient mice. After transplant, mice are monitored for wellness daily for up to 10 days. Their body weight and blood glucose (in diabetic animals) are recorded 2-3x per week. Diabetic mice are defined as have non-fasting blood glucose >250mg/dL. Mice are considered euglycemic when their non-fasting blood glucose returns to <250mg/dL. The nephrectomy procedure involves the same anesthetic regimen; renal vessels are first tied to prevent hemorrhage and kidney containing islet graft is removed.

### Histology

Islet graft and pancreas sections were fixed in 4% paraformaldehyde, embedded in optimal cutting temperature compound, and frozen on dry ice. Embedded grafts were sectioned on Leica CM3050 S (Leica Biosystems, Buffalo Grove, IL). Standard immunofluorescent staining techniques were used on 6 µm sections. Sections were blocked in 5% BSA for 30 minutes, incubated with primary antibodies overnight at 4°C, washed before incubation with secondary antibodies for 2 hours at room temperature, and finally washed again. Slide covers were secured with Hard-set Mounting Medium with DAPI (Santa Crus Biotechnology, Dallas TX). Slides were imaged on an EVOS M5000 Cell Imaging System (ThermoFisher) and color channels were merged using Fiji (http://fiji.sc/). Primary antibodies (1:100): αCD3 (17A2) and αCD45 (30-F11) were purchased from BioLegend, insulin (Catalog #: a0564) and somatostatin (Catalog #: A0566) from Dako (Carpinteria, CA). Secondary antibodies (1:1000): CF-594 and CF-488A α-Guinea Pig were purchased from MilliporeSigma (St. Louis, MO); Alexa Fluor-594 and Alexa Fluor-488 α-Rat were purchased from BioLegend.

## Acknowledgements

We thank members of the Kim group, especially Dr. R. Whitener for encouragement and advice. P.B. is a student in the Medical Scientist Training Program (MSTP) at Stanford and part of the Stanford PhD Program in Immunology. C.C. was supported by a fellowship from the Maternal & Child Health Research Institute (MCHRI) at Stanford. Work in the Kim group was supported by the JDRF Northern California Center of Excellence (to S.K.K. and M. Hebrok), NIH awards (R01 DK107507; R01 DK108817; U01 DK123743; P30 DK116074 to S.K.K), the Reid Family, H.L. Synder Foundation and Elser Trust, two anonymous donors, and the Stanford Diabetes Research Center (SDRC). Work here was also supported by the Islet Research Core in the SDRC.

